# Visual stimulation induces distinct forms of sensitization of On-Off direction-selective ganglion cell responses in the dorsal and ventral retina

**DOI:** 10.1101/2021.06.19.449131

**Authors:** Xiaolin Huang, Alan Jaehyun Kim, Héctor Acarón Ledesma, Jennifer Ding, Robert G. Smith, Wei Wei

## Abstract

Experience-dependent modulation of neuronal responses is a key attribute in sensory processing. In the mammalian retina, the On-Off direction-selective ganglion cell (On-Off DSGC) is well known for its robust direction selectivity. However, how the On-Off DSGC light responsiveness dynamically adjusts to the changing visual environment is underexplored. Here, we report that the On-Off DSGC can be transiently sensitized by prior stimuli. Notably, distinct sensitization patterns are found in dorsal and ventral DSGCs that receive visual inputs from lower and upper visual fields respectively. Although responses of both dorsal and ventral DSGCs to dark stimuli (Off responses) are sensitized, only dorsal cells show sensitization of responses to bright stimuli (On responses). Visual stimulation to the dorsal retina potentiates a sustained excitatory input from Off bipolar cells, leading to tonic depolarization of dorsal DSGCs. Such tonic depolarization propagates from the Off to the On dendritic arbor of the DSGC to sensitize its On response. We also identified a previously overlooked feature of DSGC dendritic architecture that can support direct electrotonic propagation between On and Off dendritic layers. By contrast, ventral DSGCs lack a sensitized tonic depolarization and thus do not exhibit sensitization of their On responses. Our results highlight a topographic difference in Off bipolar cell inputs underlying divergent sensitization patterns of dorsal and ventral On-Off DSGCs. Moreover, substantial crossovers between dendritic layers of On-Off DSGCs suggest an interactive dendritic algorithm for processing On and Off signals before they reach the soma.

## Introduction

Visual perception and visual neuronal responses are dynamically influenced by prior visual stimuli. Such short-term modulation is thought to underlie some visual perceptual phenomena such as saliency-based bottom-up visual attention and a rich repertoire of aftereffects and illusions^1–6^. In the early stage of the vertebrate visual system, retinal ganglion cells (RGCs) already show short-term adjustments of their responsiveness. Previous studies have mainly focused on adaptation, which refers to the decrease of sensitivity after a period of strong stimulus^7–12^. However, sensitization, the enhanced responsiveness after a strong stimulus, has been documented more recently. Studies in zebrafish, salamander, mouse and primate show that subpopulations of RGCs transiently increase their sensitivity after a period of high contrast stimulation^13–15^. That the phenomenon of RGC sensitization is conserved across species implies its functional significance. Sensitization has been proposed to complement adaptation for maintaining the responsiveness of the overall RGC population and improving the information encoding capacity and fidelity, and to contribute to the prediction of future visual inputs^13,15–17^.

The RGC population consists of diverse cell types, each conveying a distinct feature to the brain^18^. Delineating the sensitization or adaptation patterns of specific RGC types is thus necessary for a more comprehensive understanding of the retina’s neural code. In the mammalian retina, On-Off direction-selective ganglion cells (On-Off DSGC) are well-defined encoders of direction of motion, exhibiting a strong response to motion in their preferred direction and but weak response to motion in the opposite direction (null direction)^19^. However, how the light sensitivity of these cells is shaped by prior visual stimuli is not fully understood.

The On and Off responses of On-Off DSGCs are generated in different layers of their bistratified dendritic arbors, which are embedded in the On and Off sublaminae of the inner plexiform layer (IPL). The synaptic inputs onto each dendritic layer consist of glutamatergic inputs from On or Off bipolar cells, cholinergic inputs and asymmetric GABAergic inputs from On or Off starburst amacrine cells. The GABAergic inhibition is strongest when motion is in the null direction but weakest and delayed in the preferred direction, and thus plays an essential role in the direction tuning of DSGCs^20^. Although mechanisms underlying direction selectivity have been extensively studied, the adaptation or sensitization properties of the DSGC’s synaptic inputs and the resulting impacts on its spiking activity are unknown.

In this study, we address these outstanding questions by monitoring the synaptic inputs and spiking activity of On-Off DSGCs before and after a period of visual stimulation. We found that a set of iso-contrast stimuli can induce sensitization of synaptic inputs onto DSGCs and cause enhanced spiking responses. Surprisingly, we found that dorsal and ventral DSGCs exhibit distinct sensitization patterns that originate from the Off pathway. In contrast to the conventional view of segregated signal processing in the On and Off dendritic layers of the DSGC, we noted substantial dendritic crossovers between layers that may contribute to the relay of sensitization from the Off to the On pathway in the dorsal DSGC. Together, these results reveal location-dependent synaptic mechanisms underlying the divergent sensitization patterns of On-Off DSGCs receiving inputs from the upper and the lower visual fields.

## Results

### On-Off DSGC light responses can be transiently sensitized after a set of visual stimuli

To examine the influence of prior visual stimuli on the light sensitivity of On-Off DSGCs, we targeted the On-Off DSGC subtype preferring motion in the posterior direction (pDSGCs) in the Drd4-GFP transgenic mouse line^21^ for patch clamp recording. We monitored the baseline pDSGC spiking response to a 1 second flashing spot (termed “test spot”) presented every 3.5 seconds. Then, 27.5 seconds of visual stimulation (termed the “induction stimulus”) was presented to induce sensitization. We tested three types of induction stimuli at the same contrast level as the test spot: preferred direction moving spots, preferred direction drifting gratings, and contrast reversing gratings (5 repetitions of 5.5 s trials, also see Methods) (**Fig. 1a**). Immediately after the induction stimulus, the pDSGC firing rate was monitored by trials of the same test spots as those before the induction stimulus. The average firing rates of the pDSGC to the test spot before and after the induction stimulus were then used to calculate a sensitization index, which was defined as (Firing rate _After_ - Firing rate _Before_) / (Firing rate _After_ + Firing rate _Before_). Sensitized pDSGC responses are represented as positive sensitization index values, while adapted responses give negative values.

**Fig. 1.**
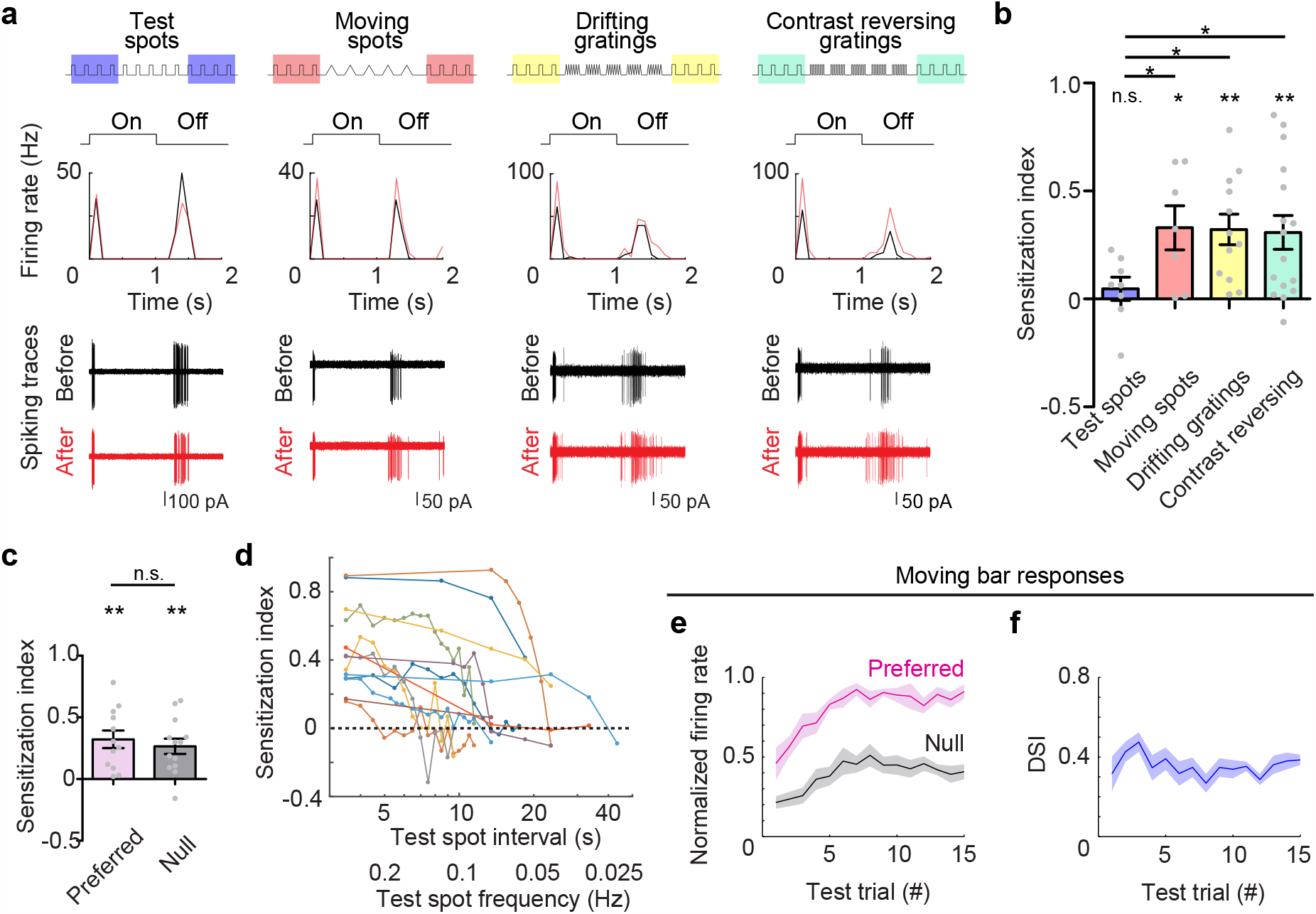
pDSGC responses are transiently sensitized after induction visual stimulation. **a**, Example pDSGC responses to 1s-duration flashing spot stimuli (“test spot”) before and after 5 repetitions of test spots, moving spots, drifting or contrast reversing gratings stimuli. Top: schematics of the stimulus protocols. Middle: Firing rates of the example cells before (black) and after (red) different stimulations. Bottom: Overlay of four repetitions of pDSGC spiking traces responding to test spots before and after different stimulations. The test spot has onset at t=0 and offset at t=1s. **b**, Summary graph comparing sensitization indices of pDSGC responses after exposure to test spots (n = 8 cells from 4 mice), moving spots in preferred direction (n = 7 cells from 3 mice), drifting gratings in preferred direction (n = 12 cells from 6 mice) or contrast reversing gratings (n = 16 cells from 6 mice). For this and subsequent plots, data with Gaussian distribution were represented as mean ± SEM, and grey dots represent individual cells. One-sample student t-test was used to test whether the sensitization index value of pDSGCs was significantly different from 0, while two-sample t-test was used for comparison between control (“test spots”) and induction visual stimulations. All p values were adjusted with FDR correction: test spots: p = 0.41; moving spots: ^*^p = 0.031; drifting gratings: **p = 0.0056; contrast reversing gratings: ^**^p = 0.0046; test spots vs moving spots: *p = 0.035; test spots vs drifting gratings: ^*^p = 0.027; test spots vs contrast reversing gratings: *p = 0.044. **c**, Comparison of pDSGC sensitization indices after 5 repetitions of drifting gratings in preferred (n = 12 cells from 6 mice) or null direction (n = 13 cells from 4 mice). Preferred direction: ^**^p = 0.0024; null direction: ^**^p = 0.0017; preferred vs null, p = 0.55. **d**, Plot of sensitization indices of individual cells with increasing interval between test spots after the induction stimulus (also see Methods) (n = 13 cells from 5 mice). Individual cells are represented in different colors. **e**, Normalized firing rate of pDSGCs in preferred and null directions relative to the maximal response of the cell in all trials. Mixed-effects analysis for repeated measurements (n = 6 cells from 3 mice): for preferred response, ^*^p = 0.014; for null response, ^*^p = 0.035. **f**, Direction selectivity index (DSI) of pDSGCs monitored over test trials. Mixed-effects analysis for repeated measurements (n = 6 cells from 3 mice): p = 0.15. See also **Supplementary Fig. 1** for spike traces showing the maintenance, extinction and repeated induction of neural sensitization in an example pDSGC.

We found that pDSGC spiking responses to test spots were significantly sensitized by all three patterns of induction stimuli. As a control, continuous presentation of test spot trials did not induce sensitization (**Figs. 1a and 1b**). For the rest of this study, we used drifting gratings as the induction stimulus to study the mechanism underlying pDSGC sensitization. Since DSGCs are direction-selective, we also tested if pDSGCs can be sensitized by drifting gratings moving in the null direction. We found that motion in both preferred and null directions can induce similar levels of sensitization in pDSGCs (**Fig. 1c**).

We next investigated the time course of the sensitization and found the following two properties. First, the sensitization is a short-term, reversible phenomenon. We were able to repeatedly induce sensitization in 82% of pDSGCs (14/17 cells) (See **Supplementary Figure 1** for an example cell). Second, the sensitization was maintained without decay as long as test spots were presented at an inter-spot interval of 3.5 seconds used in our protocol (Comparing the third trial versus the first two trials in **Supplementary Figure 1**). In our longest experiment, the pDSGC firing rate to test spots remained sensitized for 210 s. However, in the absence of continuous presentation of test spots, pDSGC light responses decayed back to the baseline level within 5-20 s after the induction of sensitization (**Fig. 1d**, see **Methods**).

Moreover, we found that the pDSGC’s response to moving stimuli can also be sensitized. We monitored the spiking activity of pDSGCs during repeated presentation of moving bar trials at the same frequency as the test spots (3.5 s each trial) in either preferred or null directions in a pseudorandom manner. We found that pDSGC firing rates in both directions show sensitization during the first 6 trials of moving bar stimulation before reaching a stable level while the direction selectivity index of the cell remains unchanged (**Fig. 1e and 1f**).

### Spiking activities of pDSGCs from the dorsal and the ventral retina show differential patterns of sensitization

Despite an overall increase of spiking activity in all pDSGCs, we noticed that pDSGCs in the dorsal and ventral regions of the retina show distinct sensitization patterns after the induction stimulus (**Figs. 2a and 2b**). We detected sensitized On responses only in the dorsal, but not in the ventral pDSGCs. The Off responses of pDSGCs were sensitized in both the dorsal and ventral groups (**Fig. 2b and 2c**). Notably, in dorsal but not ventral pDSGCs, we also observed elevated baseline spiking activity between trials of test spots, which we termed “the sustained component” (**Figs. 2b, left panel, and 2d**). In summary, for pDSGCs from the dorsal retina, both On and Off responses were sensitized by the induction stimulus, and there was a sensitized sustained component of spiking between test spot trials. However, for pDSGCs from the ventral retina, there was no sustained component and only Off responses were enhanced after the induction stimulus.

**Fig. 2.**
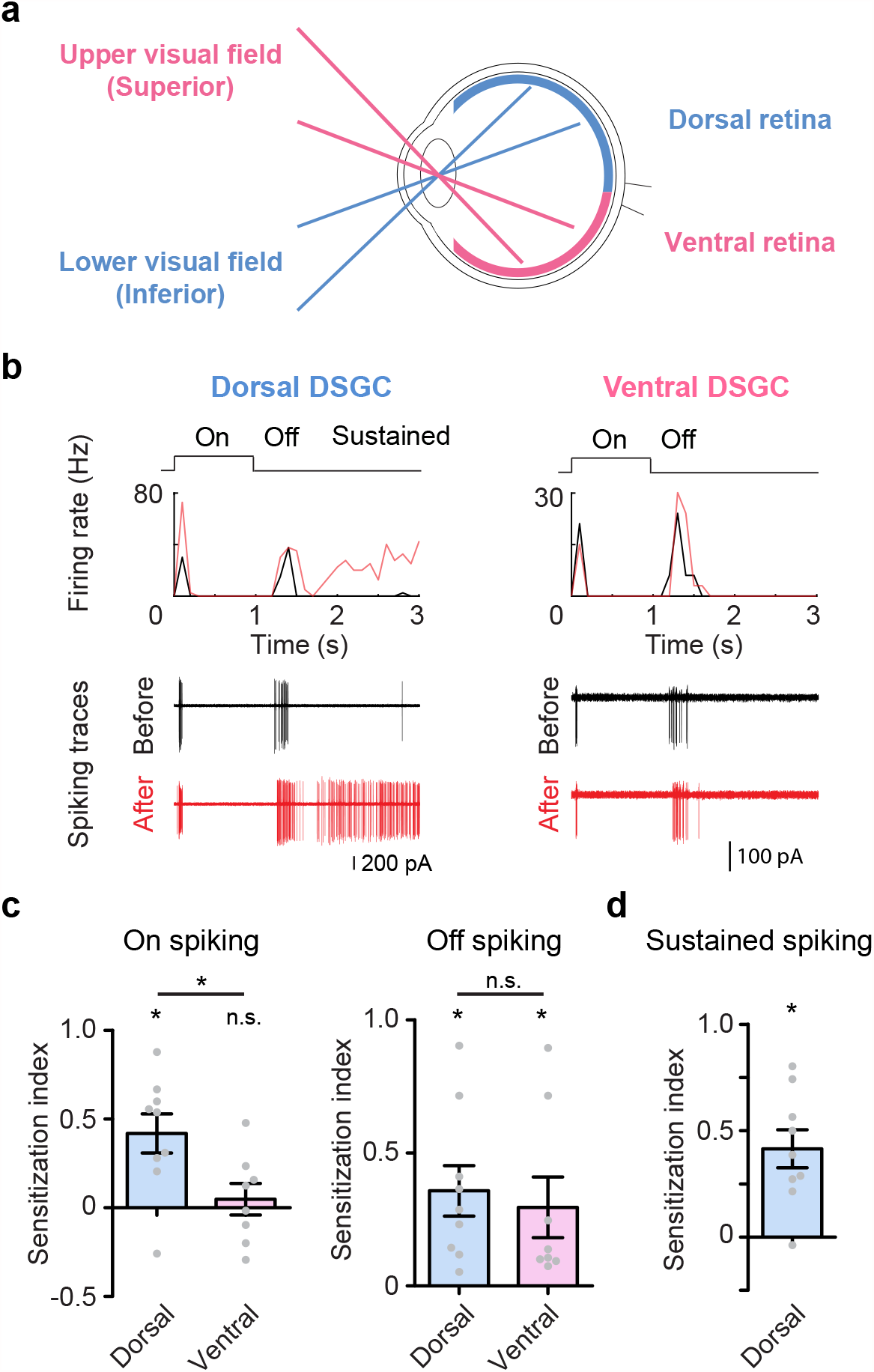
pDSGCs from the dorsal and the ventral retina show differential patterns of sensitization. **a**, Schematic diagram showing the topographic relationship of dorsal/ventral retina and the visual fields where they receive visual inputs. **b**, Firing rate plots and spiking traces of example pDSGCs from the dorsal and the ventral retina responding to test spot stimuli before (black) and after (red) the induction stimulus. **c**, Summary graphs comparing the sensitization indices of On and Off spiking between pDSGCs from the dorsal and the ventral retina. Dorsal: n = 9 cells from 3 mice; ventral: n = 8 cells from 4 mice. For On spiking: dorsal: ^*^p = 0.018; ventral: p = 0.70; dorsal vs ventral: ^*^p = 0.037. For Off spiking: dorsal: ^*^p = 0.013; ventral: ^*^p = 0.049; dorsal vs ventral: p = 0.68. **d**, Summary graph of sensitization index for the sustained component of dorsal pDSGC spiking activity. N = 9 cells from 3 mice, ^*^p = 0.011.

### Subthreshold membrane potentials of dorsal and ventral DSGCs show distinct sensitization patterns

We next examined membrane depolarization patterns that drive distinct firing patterns of dorsal and ventral pDSGCs using whole-cell current clamp recording. Spikes were digitally removed to reveal the subthreshold postsynaptic potentials (PSPs) of pDSGCs (see **Methods**). Consistent with the spiking activity, in the dorsal retina, both On and Off PSPs were sensitized after the induction stimulus, while in the ventral retina, only the Off PSPs were sensitized (**Fig. 3b**). Moreover, dorsal pDSGCs exhibit sustained depolarization of their membrane potentials between test spots after the induction stimulus (**Figs. 3a and 3c**), which corresponds to the sensitized sustained component in their spiking activities.

**Fig. 3.**
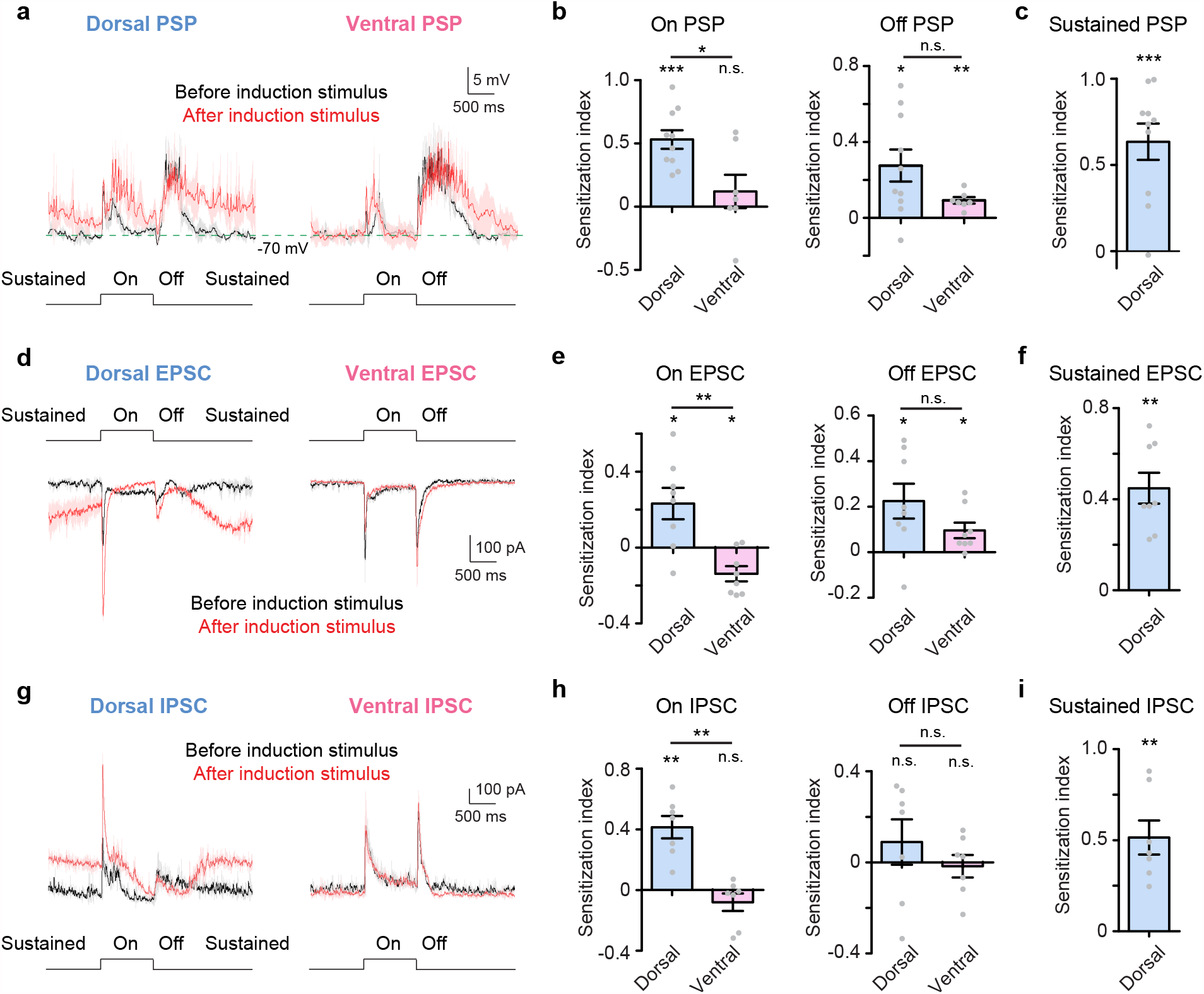
Membrane potential and synaptic currents of dorsal and ventral pDSGCs show distinct sensitization patterns. **a**, Example PSP traces of dorsal and ventral pDSGCs evoked by test spots before (black) and after (red) induction stimulus. PSP traces represent trial average (darker traces) and SEM (lighter traces). Note that in the dorsal PSP trace, there is a sustained component of elevated depolarization that persists during the time window between the test spot offset and the onset of the next test spot. **b**, Summary plots comparing the sensitization indices of PSPs between dorsal and ventral pDSGCs. Dorsal: n = 10 cells from 4 mice; ventral: n = 7 cells from 4 mice. For On PSP: dorsal: ^***^p < 0.001; ventral: p = 0.39; dorsal vs ventral: ^*^p = 0.014. For Off PSP: dorsal: ^*^p = 0.017; ventral: ^**^p = 0.0035; dorsal vs ventral: p = 0.11. **c**, Summary graph of the sensitization index of the sustained component of PSPs from dorsal pDSGCs. N = 10 cells from 4 mice, ^***^p < 0.001. **d**, Example EPSC traces of a dorsal and a ventral pDSGC evoked by test spots before (black) and after (red) induction stimulus. PSC traces represent trial average (darker traces) and SEM (lighter traces) for this and subsequent figures. Note that in the dorsal pDSGC EPSC, there is a sustained inward current that persists during the time window between the test spot offset and the onset of the next test spot. Such a sustained component was not observed in the ventral pDSGCs. **e**, Same as **b**, but for On and Off EPSCs. Dorsal: n = 8 cells from 6 mice; ventral: n = 8 cells from 5 mice. Dorsal On EPSC peak amplitude value after sensitization is relative to the elevated baseline tonic current. For On EPSC: dorsal: *p = 0.030; ventral: ^*^p = 0.026; dorsal vs ventral: ^**^p = 0.0042. For Off EPSC: dorsal: ^*^p = 0.039; ventral: ^*^p = 0.036; dorsal vs ventral: p = 0.15. **f**, Same as **c**, but for the sustained component of dorsal EPSCs. N = 8 cells from 6 mice, **p = 0.0021. **g**, Same as **d**, but for IPSCs. Note that the sustained component was also observed in the IPSC traces from the dorsal pDSGC but absent from the ventral pDSGC. **h**, Comparison of IPSC sensitization indices between dorsal and ventral pDSGCs. Dorsal: n = 7 cells from 3 mice; ventral: n = 7 cells from 2 mice. For On IPSC: dorsal: ^**^p = 0.0046; ventral: p = 0.38; dorsal vs ventral: ^**^p = 0.0014. For Off IPSC: dorsal: p = 0.47; ventral: p = 0.75; dorsal vs ventral: p = 0.50. **i**, Same as **c**, but for the sustained component of IPSCs. N = 7 cells from 3 mice, ^**^p = 0.0035.

### Synaptic inputs of dorsal and ventral DSGCs show differential patterns of sensitization

We hypothesized that the stronger depolarization of the pDSGC membrane potential after the induction stimulus may result from enhanced excitatory inputs or reduced inhibitory inputs. To determine how the synaptic inputs of pDSGC are modulated by the induction stimulus, we measured the excitatory postsynaptic currents (EPSCs) and inhibitory postsynaptic currents (IPSCs) of dorsal and ventral pDSGCs using whole cell voltage-clamp recording. After the induction stimulus, both dorsal and ventral pDSGCs showed enhanced Off EPSC responses (**Figs. 3d and 3e**). The EPSCs of dorsal pDSGCs showed an enhanced On EPSC amplitude (**Figs. 3d and 3e**), as well as an elevated sustained component between test spots (**Figs. 3d and 3f**) that corresponds to the sustained component of the spiking activity (**Figs. 2b and 2d**) and of the membrane depolarization (**Figs. 3a and 3c**).

Sensitized spiking activity was not accompanied by reduced inhibition of pDSGCs (**Figs. 3g-3i**). In dorsal pDSGCs, we detected an elevated sustained component of the IPSC after the induction stimulus similar to that of the EPSC, suggesting that the sensitization of a sustained excitatory drive to both pDSGCs and its presynaptic inhibitory neuron, likely starburst amacrine cells (SACs), which share common bipolar cell inputs^22–25^.

Taking the above results together, we found that pDSGCs transiently increased their firing after the induction stimulus. The sensitization of the spiking activity is accompanied by enhanced synaptic excitation and membrane depolarization, but not reduced inhibition. Furthermore, in the dorsal retina, sensitized pDSGCs acquired a sustained increase in their synaptic inputs, membrane potential and spiking activity between test spots.

### Synaptic activity is required for the induction of pDSGC sensitization

To investigate the mechanism underlying the induction of pDSGC sensitization, we first tested whether the membrane depolarization of the pDSGC evoked by the induction stimulus is sufficient to trigger sensitization. Instead of using the drifting grating stimulus as the induction stimulus, we mimicked drifting grating-evoked membrane potential changes in dorsal pDSGCs by directly voltage clamping the membrane potential of the pDSGC using the command voltage waveform recorded during the drifting grating stimulus (**Figs. 4a and 4b**). We found that pDSGC EPSCs were not sensitized by this direct depolarization, indicating that sensitization requires visually evoked synaptic inputs (**Figs. 4c and 4d**).

**Fig. 4.**
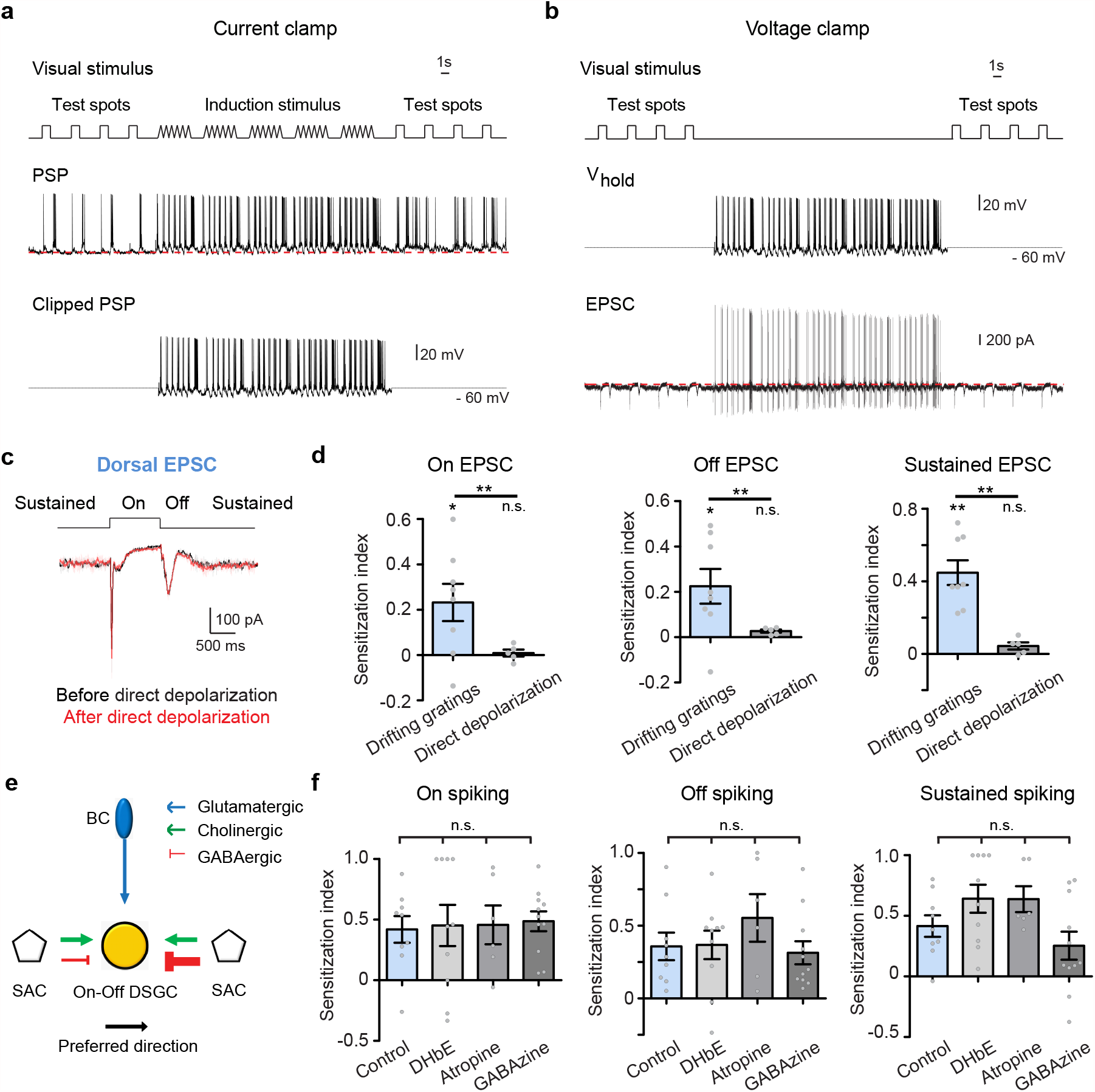
Synaptic inputs to pDSGCs are necessary for the induction of sensitization. **a**, Top: schematic shows the complete induction protocol including test spots before and after drifting gratings as the induction stimulus. Middle: Whole-cell current clamp recording of a pDSGC from the dorsal retina during the visual stimulus shown on the top. Bottom: PSP waveform evoked by drifting gratings was clipped from the PSP trace shown above. **b**, Top: schematic shows the visual stimulus protocol with only test spots but without induction stimulus (drifting gratings). Middle: Waveform of the holding potential during whole-cell voltage clamp recordings of pDSGCs. Bottom: an example EPSC trace from a dorsal pDSGC recorded with the visual stimulus protocol and the holding potential shown above. **c**, Example EPSC traces of a dorsal pDSGC during test spot stimulus before (black) and after (red) direct depolarization of the pDSGC as a replacement of drifting gratings visual stimulation. This is the same EPSC recording as the one shown in **b**. Traces are averaged from four repetitions. **d**, Comparison of the EPSC sensitization indices after drifting grating stimulus (n = 8 cells from 6 mice) versus after direct depolarization (no induction visual stimulation) (n = 5 cells from 2 mice). For On EPSC: Drifting gratings: ^*^p = 0.039; direct depolarization: p = 0.60; Drifting gratings vs direct depolarization: ^**^p = 0.0056. For Off EPSC: Drifting gratings: *p = 0.040; direct depolarization: p = 0.29; Drifting gratings vs direct depolarization: ^**^p = 0.0018. For sustained EPSC: Drifting gratings: ^**^p = 0.0014; direct depolarization: p = 0.11; Drifting gratings vs direct depolarization: ^**^p = 0.0024. **e**, Simplified schematic shows major types of synaptic inputs onto On-Off DSGCs. BC: bipolar cell; SAC: starburst amacrine cell. **f**, Comparison of the sensitization indices for pDSGC spiking in control (Ames’ solution, n = 9 cells from 3 mice) or in the presence of different receptor antagonists (DHbE: n = 10 cells from 4 mice; Atropine: n = 6 cells from 2 mice; GABAzine: n = 11 cells from 3 mice). One-way ANOVA for On spiking: p = 0.99; for Off spiking: p = 0.73; for sustained spiking: p = 0.14.

We next investigated which type(s) of synaptic signaling is required for pDSGC sensitization. A major source of synaptic inputs to pDSGCs is the SAC, which releases both acetylcholine and GABA to the DSGC (**Fig. 4e**). However, we found that pharmacological blockade of nicotinic, muscarinic, or GABA-A receptors in the retina with DHbE, atropine or gabazine respectively did not prevent the sensitization of the pDSGC spiking activity (**Fig. 4f**). This suggested that, the sensitization of the pDSGC arises from enhanced glutamate release from bipolar cells.

### Glycinergic signaling in the Off pathway contributes to sensitized glutamatergic inputs to dorsal pDSGCs

Previous studies in the vertebrate retina indicate that enhanced bipolar cell glutamate release can result from adapted presynaptic inhibition of bipolar cell terminals^14–17,26^. Since blocking GABA-A receptor signaling did not affect pDSGC sensitization (**Fig. 4f**), we next blocked another major type of presynaptic inhibition, glycinergic signaling, by bath application of strychnine while recording from dorsal pDSGCs before and after the induction stimulus. We found that glycinergic blockade significantly reduced the sensitization of dorsal pDSGC spiking activity during On and Off responses, and between test spots (**Figs. 5a and 5b**). The Off and sustained component of EPSCs also showed reduced sensitization. However, the sensitization index of the On EPSC was not affected by strychnine despite the impaired sensitization of On spiking responses (**Figs. 5c and 5d**), indicating that the sensitization of On EPSCs by itself is not sufficient to induce sensitization of On spiking responses in dorsal pDSGCs.

**Fig. 5.**
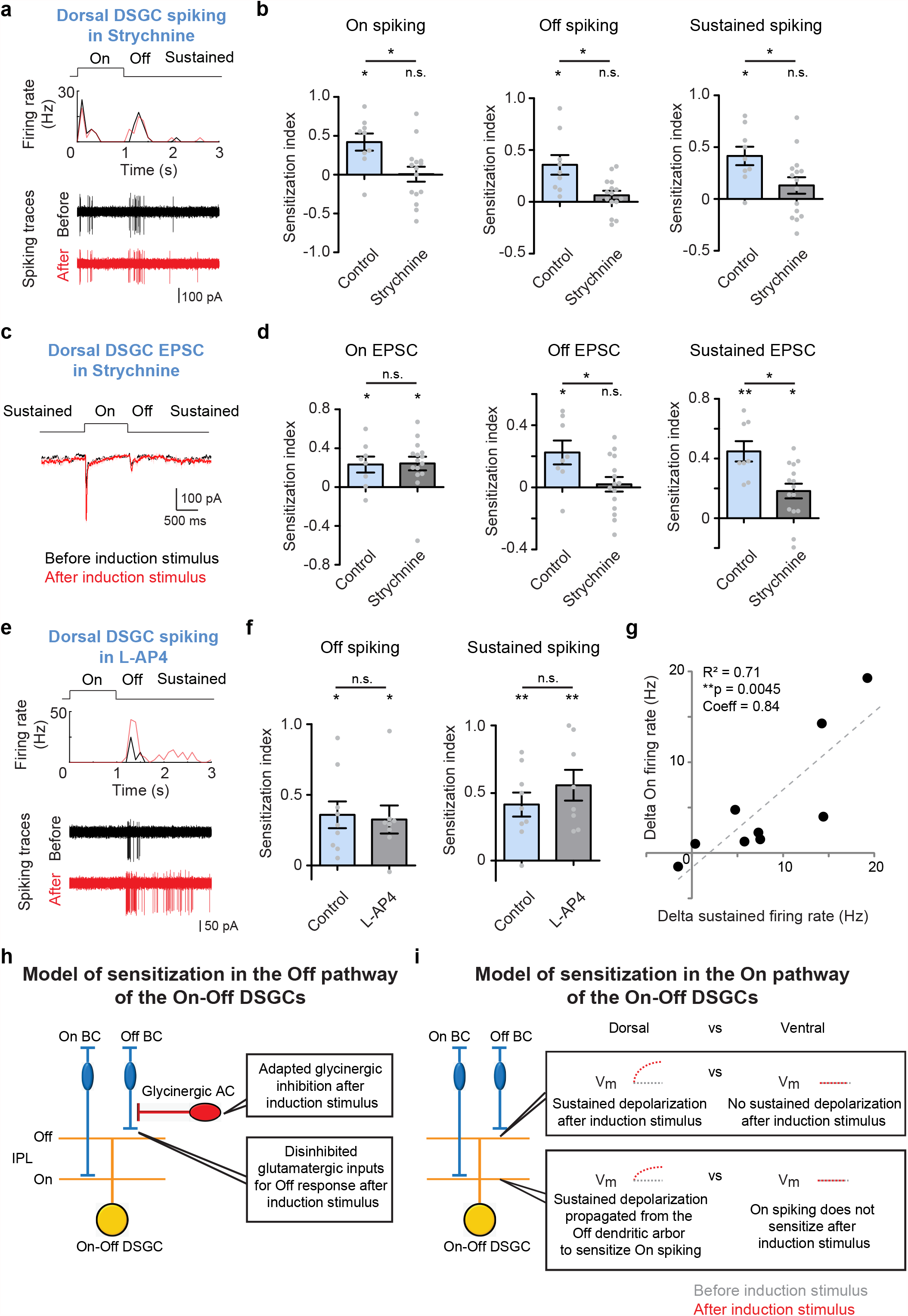
Glycinergic signaling contributes to pDSGC sensitization. **a**, Example firing rate plot and spiking traces of a dorsal pDSGC responding to test spot stimuli before and after induction stimulus in the presence of strychnine. **b**, Summary plots comparing the sensitization indices of spiking activity in control (Ames’ solution, n = 9 cells from 3 mice) and in the presence of strychnine (n = 15 cells from 5 mice). For On spiking, control: ^*^p = 0.016; strychnine: p = 0.95; control vs strychnine: ^*^p = 0.022. For Off spiking, control: ^*^p = 0.012; strychnine: p = 0.18; control vs strychnine: ^*^p = 0.018. For sustained spiking, control: ^*^p = 0.014; strychnine: p = 0.16; control vs strychnine: ^*^p = 0.047. **c**, Example EPSC traces during test spot stimulus before (black) and after (red) induction stimulus in the presence of strychnine. **d**, Comparison of the EPSC sensitization indices in control (n = 8 cells from 6 mice) versus in the presence of strychnine (n = 15 cells from 8 mice). For On EPSC, control: ^*^p = 0.033; strychnine: ^*^p = 0.011; control vs strychnine: p = 0.93. For Off EPSC, control: ^*^p = 0.040; strychnine: p = 0.76; control vs strychnine: ^*^p = 0.037. For sustained EPSC, control: ^**^p = 0.0027; strychnine: ^*^p = 0.011; control vs strychnine: ^*^p = 0.010. **e**, Example firing rate plot and spiking traces of a dorsal pDSGC responding to test spot stimuli before and after induction stimulus in the presence of L-AP4. **f**, Summary graphs comparing the sensitization indices for the Off and the sustained components of spiking activity in control (Ames’ solution, n = 9 cells from 3 mice) versus in the presence of L-AP4 (n = 8 cells from 3 mice). For Off spiking, control: ^*^p = 0.011; strychnine: ^*^p = 0.020; control vs strychnine: p = 0.81. For sustained spiking, control: ^**^p = 0.0096; strychnine: ^**^p = 0.0054; control vs strychnine: p = 0.40. **g**, Scatter plot comparing the increase of firing rates of the On spiking response versus that of the sustained component. Black dots represent individual cells (n = 9 cells from 3 mice), and dashed line indicates linear regression fit. **h and i**, A mechanistic model of sensitization in the direction-selective circuit. Schematic diagrams show side views of the laminar organization of bipolar cells (BCs), glycinergic amacrine cells (ACs) and On-Off DSGCs in the inner plexiform layer (IPL). In the Off pathway, the presynaptic glycinergic AC adapts and disinhibits Off BC after induction stimulus. Therefore, the DSGC Off responses is enhanced due to higher glutamate release from Off BCs (**h**). Moreover, dorsal DSGC also gained an elevated baseline depolarization originating from Off BCs. Such sustained depolarization propagates to On dendritic arbors and enhances the subsequent On response. In contrast, ventral DSGCs do not have sustained depolarization and thus only show sensitized Off responses but not On responses after induction stimulus (**i**).

Since a well-established role of glycinergic inhibition is to mediate crossover inhibition from the On to the Off pathway via glycinergic AII amacrine cells^27^, we tested whether On bipolar cell activity is required for the glycinergic signaling underlying pDSGC sensitization in the Off pathway. We bath applied the mGluR6 agonist L-AP4 to silence rod and On bipolar cells during visual stimulation. As expected, the On spiking response of the dorsal pDSGC was abolished in L-AP4. However, we still observed sensitized Off responses and sustained components after the induction stimulus **(Figs. 5e and 5f**). This result shows that 1) On bipolar cell activity is not involved in the sensitization of Off bipolar cell signaling, and 2) in the dorsal retina, the sustained pDSGC activity between test spots arises from the Off pathway.

Based on the above results, our working model for the sensitization in the Off pathway is that the induction stimulus triggers synaptic depression at the glycinergic synapse from amacrine cells to Off bipolar cells, which leads to increased glutamate release from Off bipolar cells to pDSGCs. Glycinergic disinhibition of Off bipolar cells causes sensitized pDSGC Off responses, as well as sustained depolarization of membrane potential between test spots in dorsal pDSGCs (**Fig. 5h**).

### Off-to-On crossover excitation within the bistratified pDSGC dendrites contributes to the sensitization of the On spiking response in the dorsal retina

In dorsal pDSGCs, blocking glycinergic signaling in the retina impaired the sensitization of the On spiking response (**Figs. 5a and 5b**), even though the On EPSC was not affected (**Figs. 5c and 5d**), suggesting an alternative glycinergic mechanism underlying the sensitization of the On spiking activity in the dorsal retina. Since the sensitization of dorsal On spiking responses is associated with the presence of the sensitized sustained components, both of which are dependent on glycinergic signaling, we hypothesized that this sustained component between test spot trials tonically increases the excitability of dorsal pDSGCs to boost their On spiking responses. We reasoned that if the sustained depolarization of the pDSGC is important for the sensitization of its On spiking response, we would expect to see a positive correlation between the two. We calculated the correlation coefficient between the change of sustained firing rate and that of the On firing rate, and indeed found a strong correlation between these two components (**Fig. 5g**, R^2^ = 0.71, **p = 0.0045, Coeff = 0.84). These experimental results support an important role of the sustained depolarization of the dorsal pDSGC after the induction stimulus in the sensitization of its On response.

How did the sustained depolarization originating from Off bipolar cells influence the On response of the pDSGC? One route is the electrotonic spread of the depolarization in the pDSGC Off dendritic arbor through the soma to the On dendritic arbor (**Fig. 6a, left**). Interestingly, we noted an alternative route for the Off-On crosstalk within the bistratified pDSGC dendritic morphology (**Fig. 6a, right**). By two-photon imaging of dye-filled pDSGCs, we noted frequent crossovers of dendritic branches from one dendritic layer to the other. On average, about 30% of the total dendritic arbors of a pDSGC originate from the other layer through crossover dendrites (**Fig. 6b**, 29.5 ± 3.2 % of the total dendritic length, mean ± SEM, n = 21 cells). The majority of the crossover segments branched from the On layer into the Off layer (the red dendrites in **Figs. 6b and 6c**). And the majority of the crossover dendrites started diving from one layer to the other at a radial distance of around 40-80 μm away from the soma (**Figs. 6d and 6e**), which mainly falls in the distal half of the pDSGC dendritic field radius (105.3 ± 2.0 μm, n = 21 cells).

**Fig. 6.**
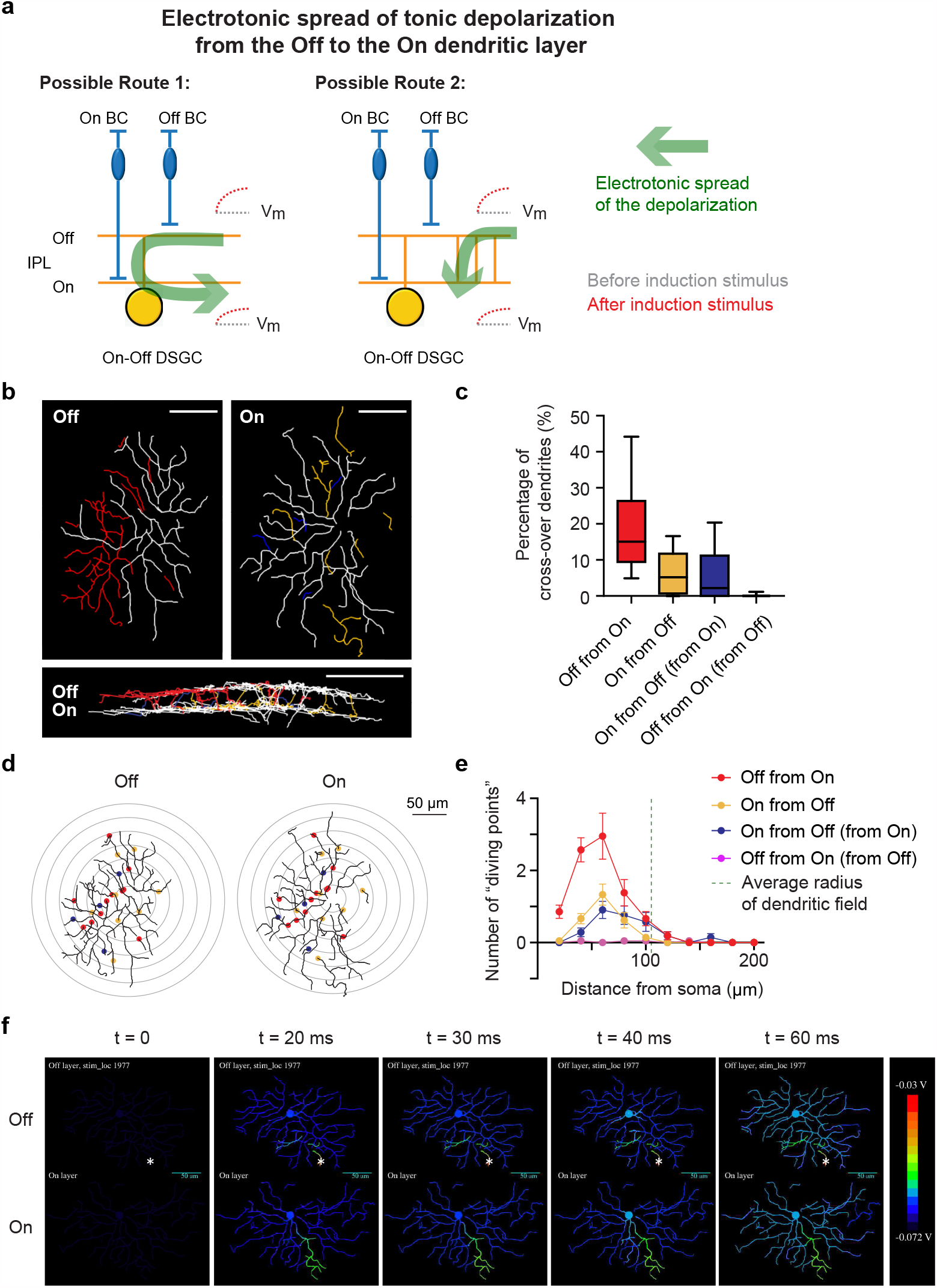
pDSGC dendrites show extensive crossovers between On and Off layers. **a**, Schematics showing two possible routes for the electrotonic spread of depolarization from the Off to the On dendritic layers of the bistratified On-Off DSGC. **b**, Dendritic morphology of an example pDSGC, including dendrites staying in one single layer (white), On dendrites crossing over to the Off layer (red), Off dendrites crossing over to the On layer (yellow), and dendrites from On to Off and then back to On layer (blue). **c**, Summary box plots representing the percentages of dendritic crossovers over total dendritic lengths. Data were represented as median ± IQR (n = 21 cells). **d**, The same example cell as **Fig. 6b**, but with labels showing the locations of the starting points where dendrites started to cross from one layer to the other (“the diving points”). The colors were coded as **Fig. 6b-6c**. Concentric rings represent radial distance from soma. **e**, Sholl analysis of the diving points (n = 21 cells). **f**, A heat map showing the membrane potentials of the pDSGC Off and On layers in response to a simulated bipolar cell input onto a location (indicated by *) in the Off layer. Model parameters: Ri = 100 Ohm-cm, dendritic dia factor = 0.5. See also **Supplementary Fig. 2** for more stimulation locations and **Supplementary Fig. 3** for brackets of parameters used in the simulation model.

To assess the functional importance of dendritic crossover in signal propagation between On and Off pathways, we simulated the electronic spread from one dendritic layer to the other in a detailed biophysical model of the pDSGC based on the reconstruction of a representative dye-filled cell (**Fig. 6f**). We simulated a single synaptic input from a presynaptic compartment that represented a bipolar cell voltage-clamped with a pulse of 100 ms duration. This input was placed at different locations throughout the Off dendritic arbor of the pDSGC while the membrane potential changes in both the On and Off dendritic layers were monitored. We found that dendritic crossovers provide shortcuts for fast and efficient spread of depolarization from the Off layer to the On layer bypassing the soma (**Fig. 6f**). Therefore, the direct route through dendritic crossovers and the transomatic route together provide plausible physical substrates for the cross-layer influence of sustained Off bipolar cell inputs on the On responses of dorsal pDSGCs during sensitization.

Based on the above experimental and modeling results, our working model for the sensitized pDSGC On response in the dorsal retina is that after the induction stimulus, dorsal pDSGCs receive enhanced tonic glutamatergic inputs from Off bipolar cells, which depolarize the membrane potential and propagate to the On dendritic layers of pDSGCs, increasing the excitability of the cell. As a result, the subsequent On stimulus triggers a stronger On response (**Fig. 5i, dorsal**). In contrast, ventral pDSGCs lack the sustained depolarization after the induction stimulus, and therefore did not exhibit sensitized On responses (**Fig. 5i, ventral**).

### Development of neural sensitization in pDSGCs

Since the sensitization of the pDSGC depends on the glycinergic circuitry that shapes the Off bipolar cell activity, we postulated that the development of the sensitization should coincide with the period when bipolar cell connectivity matures. Previous studies in rodents have shown that the integration of Off bipolar cells into the retinal network starts at around postnatal day 8 (P8) and continues for several weeks after the eye opening at P14^28–34^. We did not detect sensitization of either On or Off spiking responses of dorsal pDSGCs at the early stage of bipolar cell innervation at P12-13, nor did we detect the sustained component in dorsal pDSGCs at this stage (**Fig. 7a and 7c**). Therefore, the emergence of the sensitization and the sustained component of pDSGC occurs after eye opening, which overlaps with the maturation timeline of both glycinergic inhibition and bipolar cell connectivity in the rodent retina^28,30^.

**Fig. 7.**
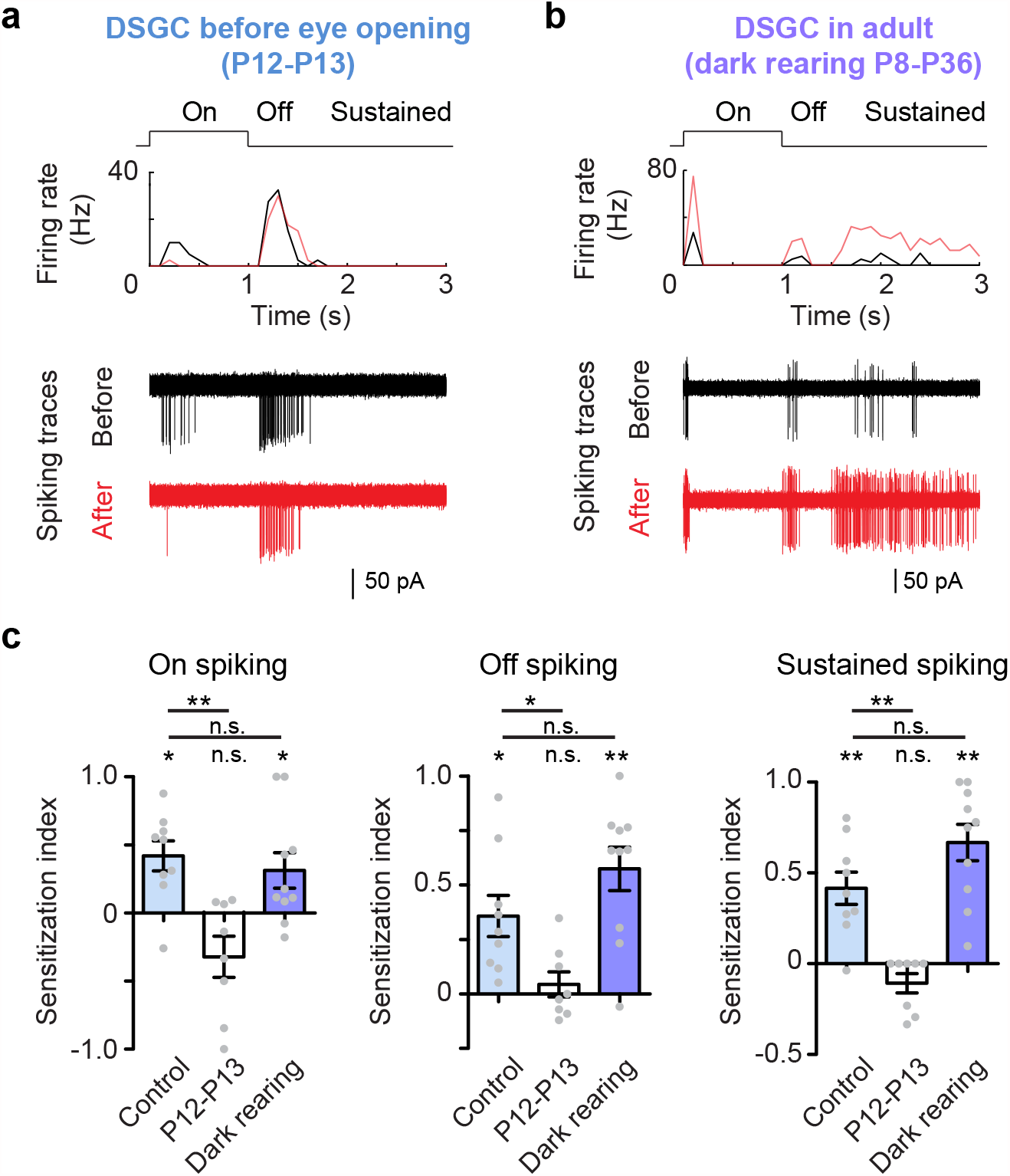
Sensitization of pDSGC light responses develops after eye opening and persists with dark rearing. **a**, Example firing rate plot and spiking traces of a dorsal pDSGC responding to test spots before and after induction stimulus from a mouse before eye opening at P12. **b**, Example firing rate plot and spiking traces of a dorsal pDSGC responding to test spots from an adult mouse dark reared during P8-P36. **c**, Summary graphs comparing the sensitization indices of dorsal pDSGC spiking from control mice (n = 9 cells from 3 mice), P12-P13 mice (n = 8 cells from 2 mice) and dark-reared adult mice (n = 10 cells from 4 mice). All p values shown here were adjusted with FDR correction. For On spiking, control: ^*^p = 0.013; P12-P13: p = 0.11; dark rearing: ^*^p = 0.050; control vs P12-P13: ^**^p = 0.0041; control vs dark rearing: p = 0.54. For Off spiking, control: ^*^p = 0.012; P12-P13: p = 0.50; dark rearing: ^**^p = 0.0015; control vs P12-P13: ^*^p = 0.028; control vs dark rearing: p = 0.16. For sustained component: control: ^**^p = 0.0048; P12-P13: p = 0.10; dark rearing: ^**^p = 0.0014; control vs P12-P13: ^**^p = 0.0015; control vs dark rearing: p = 0.11.

We next asked whether the visual experience after eye-opening is required for the development of pDSGC sensitization. We reared mice in dark from P8 to P36 and then compared the sensitization indices of pDSGCs from these mice to those of the controls. Dark rearing did not alter the normal pattern of sensitization: dorsal cells still exhibited sustained elevation of baseline firing and enhanced light responses to test spots after the induction stimulus (**Fig. 7b and 7c**). Therefore, the sensitization of pDSGCs developed after eye opening but was independent of visual experience.

### Sensitization of other types of RGCs in the mouse retina

The sensitized Off bipolar cell inputs detected in our study may influence multiple postsynaptic targets in addition to pDSGCs. To test if other RGC types also receive sensitized Off bipolar cell inputs in the dorsal retina, we focused on alpha ganglion cells, which can be conveniently targeted by their large soma sizes for recording. On transient (tOn), On sustained (sOn), Off transient (tOff) and Off sustained (sOff) alpha cells were identified based on their large soma sizes and typical light responses as reported previously^35^. We noted a subset of RGCs with large somas had sustained Off responses but exhibited biphasic Off spiking activity (**Supplementary Figure 2a**, n = 7 cells from 5 mice). Here we tentatively classify them as sOff alpha cells. We found that these four types of alpha cells had similar levels of peak firing rate (Kruskal-Wallis test, p = 0.39) but different baseline firing rates (Kruskal-Wallis test, *p = 0.040, **Supplementary Figure 2b**), which agrees with previous descriptions^35^.

We then calculated the sensitization index of alpha cells, and found that sOff alpha cells also showed sensitization after the induction stimulus (**Fig. 8a**). We noted that the subset of sOff alpha cells with biphasic Off responses were located exclusively in the dorsal retina (7 out of 8 dorsal cells, 0 out of 4 ventral cells, **Fig. 8c**). Moreover, only the dorsal biphasic sOff alpha cells showed sensitized responses to test spots after the induction stimulus (**Figs. 8b and 8c**). Sensitization of biphasic Off responses in both pDSGCs and sOff alpha cells in the dorsal retina supports our working model of sensitized Off bipolar cell sustained signaling in this region.

**Fig. 8.**
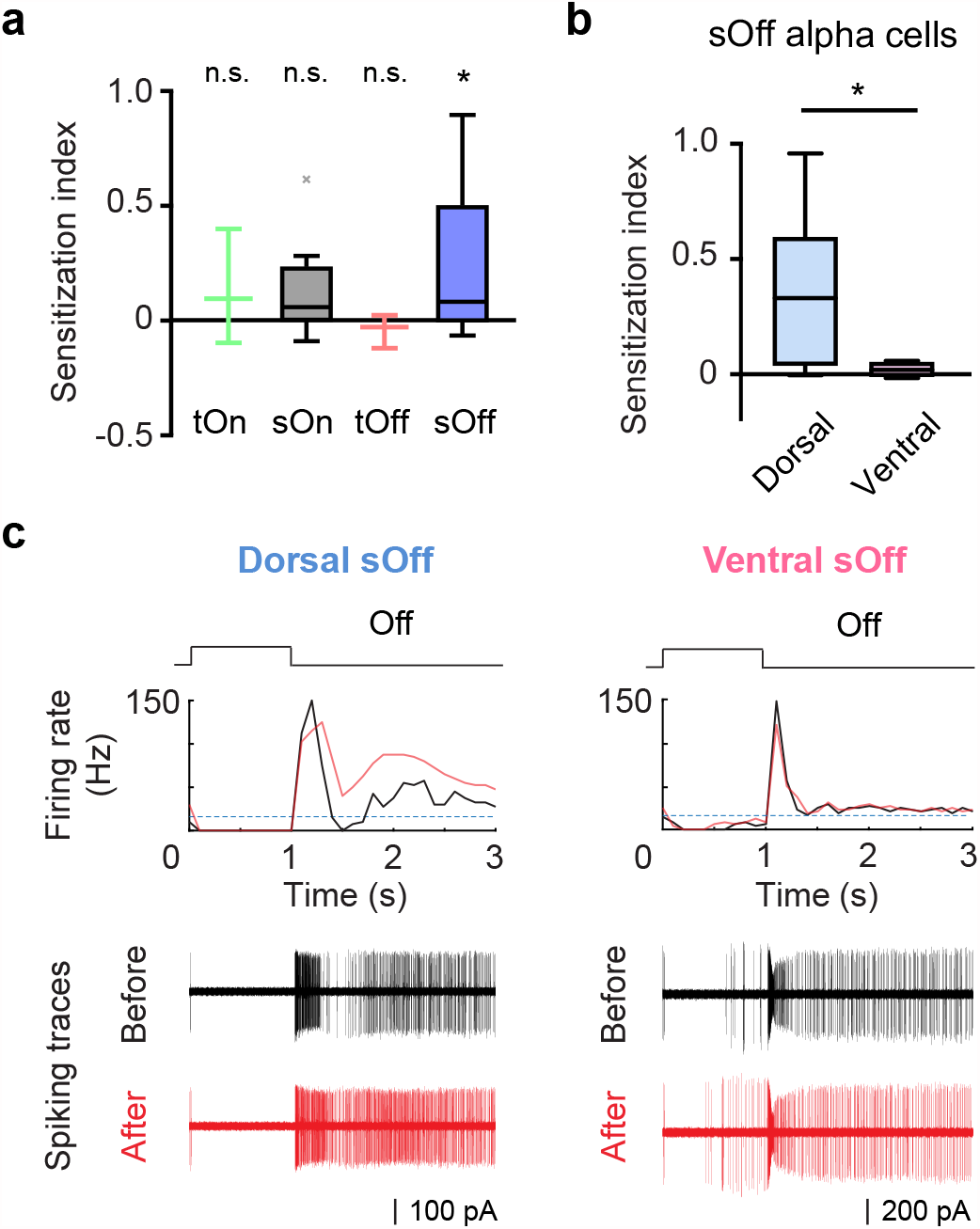
Sensitization is detected in sustained Off alpha ganglion cells. **a**, Summary box plot of sensitization indices for four types of alpha ganglion cells. Single sample Kolmogorov-Smirnov test: tOn, n = 3 cells from 3 mice, p = 0.75; sOn, n = 11 cells from 9 mice, p = 0.084; tOff, n = 3 cells from 3 mice, p = 0.66; sOff, n =12 cells from 8 mice, *p = 0.037. **b**, Comparison of the sensitization indices between dorsal (n = 8 cells from 5 mice) and ventral (n = 4 cells from 3 mice) sOff alpha cells. Two-sample Kolmogorov-Smirnov test, ^*^p = 0.048. **c**, Example firing rate plots and spiking traces of sOff alpha cells from the dorsal and the ventral retina during 1s-duration test spot stimuli before (black) and after (red) induction stimulus. See also **Supplementary Fig. 4** for firing patterns of alpha ganglion cells.

## Discussion

Our finding that the pDSGC responsiveness can be transiently and reversibly sensitized by visual stimulation adds to the accumulating evidence on the contextual modulation of the DSGC response in addition to its robust direction selectivity^36–40^. Notably, dorsal and ventral pDSGCs differ in their sensitization patterns. When sensitization is induced in dorsal pDSGCs, a tonic depolarization originating from Off bipolar cells onto the DSGC Off dendrites readily spreads into the On dendritic layer, causes a prolonged increase of excitability, and boosts the subsequent On spiking responses. In contrast, ventral pDSGCs lack such a sustained elevation of dendritic excitability after induction and therefore are not subject to the relay of sensitization from the Off to the On pathway. Because this dorsal-ventral difference arises from Off bipolar cell modulation, other RGC types that share common Off bipolar cell inputs with pDSGCs may have a similar divergence of sensitization patterns between the dorsal and the ventral retinal regions. Indeed, we found a comparable dorsal-ventral difference in the sensitization pattern of sOff alpha cells, which share common inputs with On-Off DSGCs from type 2 Off bipolar cells (CBC2)^22,23^.

Different adaptive properties of pDSGC responsiveness in the dorsal and the ventral retina highlight the topographic variations in the retinal code. In the mouse retina, dorsal-ventral asymmetry has been reported at multiple stages of visual processing from photoreceptor spectral sensitivity to ganglion cell sizes, densities and receptive field properties^41–44^. These specializations on the vertical axis are thought to reflect the adaptation of the retinal circuitry to the different environments in the animal’s upper and lower visual fields. However, in the direction-selective circuit, while extensive studies have focused on the robustness of direction selectivity across the retina under diverse visual conditions, regional differences of the circuit on the vertical axis are underexplored. A decrease in On-Off DSGC dendritic field size from the dorsal to the ventral retina has been reported^45^. Moreover, On starburst amacrine cells in the dorsal retina can reverse their contrast polarity under certain visual stimulation conditions, a phenomenon that likely originates from region-specific photoreceptor properties and may contribute to the switch of the DSGC directional preference^36,37,46^.

In this study, we found that Off bipolar cell signaling in the dorsal and the ventral retina is differentially modulated by visual experience, which contributes to the distinct sensitization patterns of postsynaptic RGC targets including On-Off pDSGCs. An induction stimulus triggers elevated baseline firing and enhanced On and Off responses in dorsal pDSGCs, but only a transient increase of Off responses in ventral cells. Therefore, dorsal and ventral pDSGCs report the changing visual scenes differently to their downstream targets in the brain including the superficial layer of the superior colliculus and the shell region of the dLGN^21,47–49^. As an interesting parallel, the dorsal retina receives more balanced On and Off stimuli on the ground while the ventral retina receives predominant Off stimuli in the sky^50^. These observations suggest that differential processing strategies of dorsal and ventral retinal circuits may underlie different coding principles of upper and lower visual field information, and help serve the animal’s behavioral demands in its ecological niche.

Previous studies on RGC sensitization primarily focused on contrast adaptation, a condition under which induction stimuli have a higher contrast than the test stimulus^13–17^. In this study, we found that the sensitization of pDSGCs can be induced by other forms of visual stimulation, such as moving spots, drifting and contrast-reversing gratings, at the same contrast as the testing stimulus.

Despite different forms of induction stimuli, our results and other studies^13–17^ indicate that RGC sensitization in several species involves short-term disinhibition of bipolar cells. This common mechanistic origin implies that the phenomenon of sensitization is not bound by the category of the induction stimulus *per se*, but reflects the synaptic plasticity rules that permit short-term modulation of bipolar cell signaling under multiple stimulus conditions.

In the dorsal retina, the tonic elevation of Off bipolar cell inputs after sensitization permits a specific mode of crossover signaling from the Off to the On dendritic layers of the bistratified pDSGC. The electrotonic spread of sustained depolarization from the Off to the On dendritic layers depends on the dendritic architecture and membrane properties. In this context, the dendritic crossovers between the On and the Off layers of On-Off DSGCs, which are evident in published retinal studies^47,51^ but have not been investigated, are particularly relevant and caught our attention. Here, we provide the first quantification of this dendritic feature in mouse pDSGCs. We found that dendritic crossover is present in every pDSGC. For a given cell, a significant fraction of dendrites in one layer originates from the other layer. These direct connections between the On and the Off dendritic layers bypassing the soma indicate a more direct route for membrane depolarization to spread across dendritic layers. Therefore, elevated membrane excitability in the Off dendritic layer can be more readily relayed to the On dendritic layer to sensitize its On response. Interestingly, studies in the rabbit retina have demonstrated that spikes of On-Off DSGCs are initiated in the dendritic arbors^52,53^. In this context, dendritic crossovers may significantly influence local spike initiation at the On layer upon sensitization. Future experimental and modeling studies will provide more insights into functional implications of On-Off DSGC dendritic crossovers during visual processing.

## Acknowledgements

We thank Chen Zhang for managing the mouse colony. This work was supported by NIH R01 EY02416, R01 NS109990 and the McKnight Scholarship Award to W.W., NSF GRFP DGE-1746045 to J.D., NIH F31 EY029156 to H.E.A., and NIH EY022070 to RGS. The authors declare no competing financial interests.

## Author contributions

X.H. and W.W. conceived the concept, designed the experiments and wrote the paper. X.H. conducted all the physiological experiments and data analysis. A.J.K. conducted morphological analysis of pDSGCs in Fig. 6 and loose cell-attached recording of alpha cells in Fig. 8. H.E.A. conducted supporting experiments (data not shown). J.D. collected the dye-filled pDSGC morphology dataset in Fig. 6. R.G.S. conducted the computational modeling in Fig. 6 and helped edit the paper.

## Methods

### Animals

*Drd4-GFP* mice of ages P12-13 or P22-P53 of either sex were used in this study to label On-Off DSGCs that prefer motion in the posterior direction (pDSGCs). This mouse line was originally developed by MMRRC (http://www.mmrrc.org/strains/231/0231.html) in the Swiss Webster background and subsequently backcrossed to C57BL/6 background. All procedures for mouse maintenance and use were in accordance with the University of Chicago Institutional Animal Care and Use Committee (Protocol number ACUP 72247) and in conformance with the NIH Guide for the Care and Use of Laboratory Animals and the Public Health Service Policy.

### Whole-mount retina preparation

Mice were dark adapted for more than 30 min, anesthetized with isoflurane and then euthanized by decapitation. Under infrared light, retinas were isolated from the pigment epithelium layers and cut into halves at room temperature in Ames’ medium (Sigma-Aldrich, St. Louis, MO) bubbled with 95% O_2_/5% CO_2_. The retinas were then mounted with ganglion-cell-layer up on top of a ∼1.5 mm^2^ hole in a small piece of filter paper (Millipore, Billerica, MA). Cells in the center of the hole were used for experiments.

### Visual stimulation

A white organic light-emitting display (OLEDXL, eMagin, Bellevue, WA; 800 × 600 pixel resolution, 60 Hz refresh rate) was controlled by an Intel Core Duo computer with a Windows 7 operating system and presented to the retina at a resolution of 1.1 μm/pixel. To be noted, the light spectrum of the OLED does not cover the absorption spectrum of S opsin and thus only activates rhodopsin and M opsins^54,55^. In this context, our light stimuli evoked DSGC EPSCs with comparable amplitudes in the dorsal and the ventral regions (data not shown), consistent with the even distribution of M-opsin expressing cones on the vertical axis^55,56^. All visual stimuli were generated using MATLAB and Psychophysics Toolbox^57^, projected through the condenser lens of the two-photon microscope focused on the photoreceptor layer, and centered on the neuron somas.

### Two-photon guided electrophysiology recording

Retinas were perfused with oxygenated Ames’ medium with a bath temperature of 32–34°C. GFP-labelled pDSGCs in *Drd4-GFP* mice were targeted using a two-photon microscopy (Scientifica) and a Ti:sapphire laser (Spectra-Physics) tuned to 920 nm. Data were acquired using PCLAMP 10 software, Digidata 1550A digitizer and MultiClamp 700B amplifier (Molecular Devices, Sunnyvale, CA), low-pass filtered at 4 kHz and digitized at 10 kHz.

For loose cell-attached recordings, electrodes of 3.5–5 MΩ were filled with Ames’ medium. For current-clamp whole cell recording (I = 0), electrodes were filled with a potassium-based internal solution containing 120 mM KMeSO_4_, 10 mM KCl, 0.07 mM CaCl_2_·2H_2_O, 0.1 mM EGTA, 2 mM adenosine 5′-triphosphate (magnesium salt), 0.4 mM guanosine 5′-triphosphate (trisodium salt), 10 mM HEPES, 10 mM phosphocreatine (disodium salt), pH 7.25. For voltage-clamp whole cell recording, electrodes were filled with a cesium-based internal solution containing 110 mM CsMeSO_4_, 2.8 mM NaCl, 5 mM TEA-Cl, 4 mM EGTA, 4 mM adenosine 5′-triphosphate (magnesium salt), 0.3 mM guanosine 5′-triphosphate (trisodium salt), 20 mM HEPES, 10 mM phosphocreatine (disodium salt), 5 mM N-Ethyllidocaine chloride (QX314) (Sigma), pH 7.25.

Light-evoked EPSCs and IPSCs of pDSGCs were isolated by holding the cells at reversal potentials (0 mV for GABAergic and −60 mV for cholinergic). Liquid junction potential (∼10 mV) was corrected. In **Fig. 4**, to mimic the pDSGC activation pattern during drifting grating stimulus, we selected representative current-clamp recordings of pDSGC membrane potential waveforms during drifting gratings, and used them as command potential waveforms in voltage-clamp experiments to replace the visual induction stimulus.

To investigate the contributions of different types of synaptic transmission to pDSGC sensitization, a synaptic agonist or antagonist was included in the Ames’ medium: 0.008 mM Dihydro-b-erythroidine hydrobromide (DHbE; Tocris) for blocking nicotinic cholinergic receptors; 0.002 mM Atropine (Sigma) for blocking muscarinic cholinergic receptors; 0.0125 mM GABAzine (SR-95531, Tocris) for blocking GABA-A receptors; 0.001 mM Strychnine (Sigma) for blocking glycinergic receptors; 0.005 mM L-AP4 (Tocris) for activating type 6 metabotropic glutamatergic receptors (mGluR6) and blocking the On signaling pathways.

### Analysis of electrophysiological data

For the measurement of the baseline light responsiveness, 5 repetitions of test spots were presented. Responses during the second to the fifth test spots were averaged as baseline light responses (N_Before_) and the response during the first test spot was discarded to avoid the impact of the fast adaptation after the onset of visual stimulus from the long-term dark adaptation^58^. 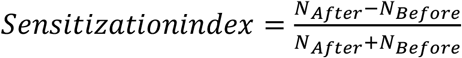 was used to quantify the strength of sensitization, where N is the averaged pDSGC responses to the four test spots right before (N_Before_) and after (N_After_) the period of stationary flash spots (same as test spots), moving spots, drifting gratings or contrast reversing gratings stimuli. A higher positive sensitization index value indicates stronger sensitization, while a negative sensitization index value indicates adaptation. N is firing rate for loose cell-attached recording data, subthreshold integral area for PSP and peak amplitude for PSC.

The time windows used to separate On, Off and sustained components were determined by the EPSC waveforms of dorsal pDSGCs which had three clear peaks. The mean of the boundary between the Off and the sustained components was ∼700 ms (n = 8 cells from 6 mice). Defining the onset of the test spot as t = 0, the On response time window was 0 – 1s; the Off response time window was 1 – 1.7 s; and the sustained component time window was 1.7 – 3 s. The same time windows were used for analyzing spiking, PSP and PSC data of both dorsal and ventral pDSGCs.

Data were analyzed using PCLAMP 10, MATLAB and GraphPad Prism. For whole-cell patch clamp recordings, membrane tests were performed to check the recording quality, and recordings with series resistances > 25 MΩ or a ratio of input resistance to series resistance < 10 were discarded.

### Dendritic Tracing

GFP-labelled pDSGCs in *Drd4-GFP* mice were targeted using a two-photon microscopy and filled with 25 μM Alexa Fluo 594 (Life Technologies). DSGC dendrites were traced from z-stacked images in ImageJ using the open source software Simple Neurite Tracer (SNT). On and Off layers were identified and separated using NeuronStudio, and then dendritic length and dendritic arbor diameter^45^ were calculated in MATLAB.

Two criteria were used to determine a dendritic segment as a crossover dendrite originating from one layer into the other: 1) The dendrite had at least 5 microns of segments remaining in the original layer before diving down. 2) The dendrite crossed over the gap between two layers and stratified into the other layer. The crossover dendrites were then classified into four subtypes labelled in different colors (**Fig. 6**). Red: Off dendrites originated from On dendrites (“Off from On”); Yellow: On dendrites originated from Off dendrites (“On from Off”); Blue: On dendrites originated from the Red “Off from On” crossover dendrites (“On from Off (from On)”); Magenta: Off dendrites originated from the Yellow “On from Off” crossover dendrites (“Off from On (from Off)”).

### Computational simulation

A model of the pDSGC was developed from a real ganglion cell morphology (the cell in Fig. 6b) that had been reconstructed from 2-photon images. The dendritic diameters were adjusted by multiplying by a constant termed the “dendritic dia factor” (0.3 - 0.8; typically 0.5) to correct for enlargement of dendrite diameter during imaging. The morphology was discretized into a compartmental model (compartment size = 0.01 lambda, ∼2200 compartments; Ri=100 Ohm-cm, Rm=20,000 Ohm-cm^2^, Vrev= −70mV) without voltage-gated ion channels to simulate subthreshold behavior in the pDSGC. The pDSGC model was stimulated at one dendritic location, and simultaneously the evoked membrane voltages were recorded at another set of locations. The stimulus was a single synaptic input from a presynaptic compartment that represented a bipolar cell voltage-clamped with a pulse of 100 ms duration. The postsynaptic conductance in the pDSGC was 2000 pS, with a reversal potential of 0 mV. Movies were generated by displaying the morphology as 2 separate images, each showing one of the pDSGC’s dendritic arborization layers. The other layer in each image was made transparent. To show the spread of depolarization through the dendritic arbor, the dendritic membrane voltage was displayed as a heat map, with violet representing −72 mV and red representing −30 mV. The movie frame interval was 1 ms. The model simulations and movies were constructed with the simulation language Neuron-C^59^.

### Statistical analysis

Grouped data with Gaussian distribution were presented as mean ± SEM in summary graphs with scattered dots representing individual cells. Two-sided one-sample t-test was performed to test whether the sensitization index value was significantly different from 0, while Two-sided two-sample t-test was used to compare two sample groups. Grouped data with non-Gaussian distribution were presented as median ± IQR in box plots, and Kolmogorov-Smirnov test was applied. For multiple comparisons, p values were adjusted with false discovery rate (FDR) correction^60^. P < 0.05 was considered significant; n.s. stands for no significance; *p < 0.05; **p < 0.01; ***p < 0.001. The number of experimental repeats were indicated in figure legends.

## Data and Code Availability

All relevant data collected and analyzed in this study are available from the authors on reasonable request. The Neuron-C simulation package and codes for the pDSGC model are available at ftp://retina.anatomy.upenn.edu/pub/nc.tgz.

