## Supplemental information for "Visual stimulation induces distinct forms of sensitization of On-Off direction-selective ganglion cell responses in the dorsal and ventral retina"

#### Supplementary Figure 1

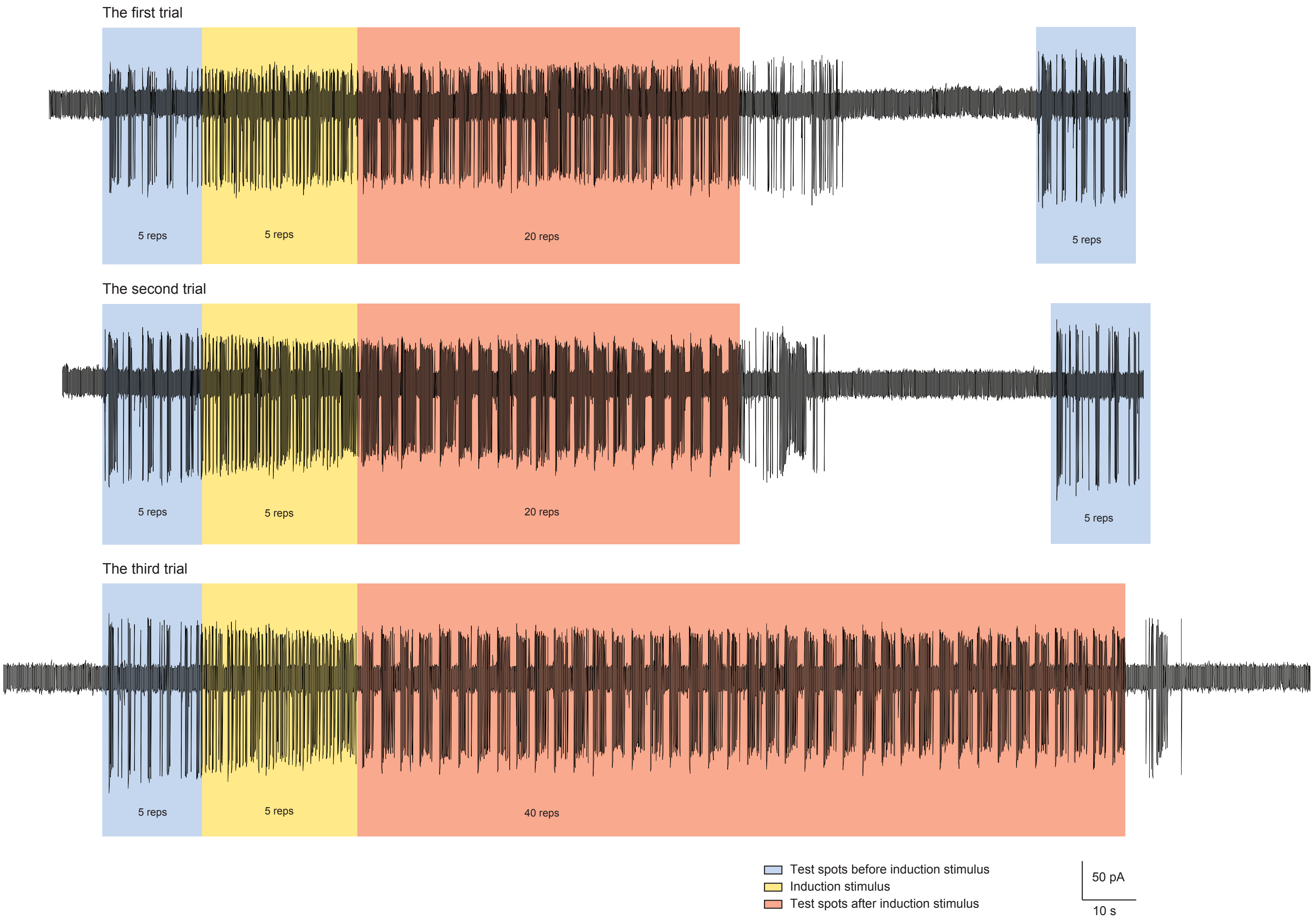


**Supplementary Figure 1. Example spiking traces of a pDSGC represent the maintenance, extinction, and repeated induction of neural sensitization. Related to Fig. 1.**

Upper, middle and lower traces represent pDSGC spiking responses during the first, second and third trials of the induction protocol.

### Supplementary Figure 2

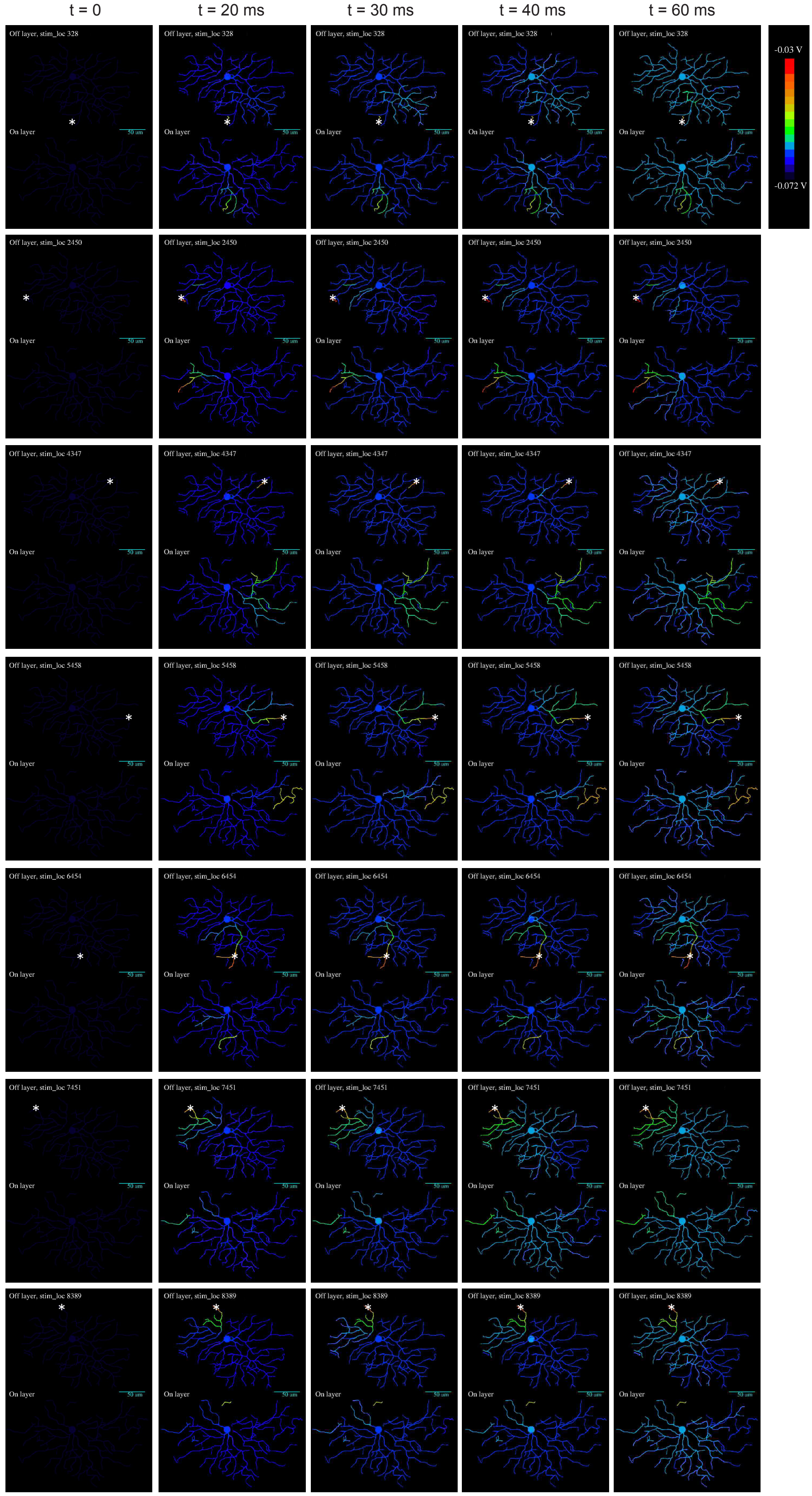


**Supplementary Figure 2. Heat maps of pDSGC membrane potentials after stimulation at different locations in the Off layer. Related to Fig. 6.**

The stimulated locations are indicated by * in the heat map plots. For these simulation videos, Ri = 100 Ohm-cm, dendritic dia factor = 0.5.

#### Supplementary Figure 3

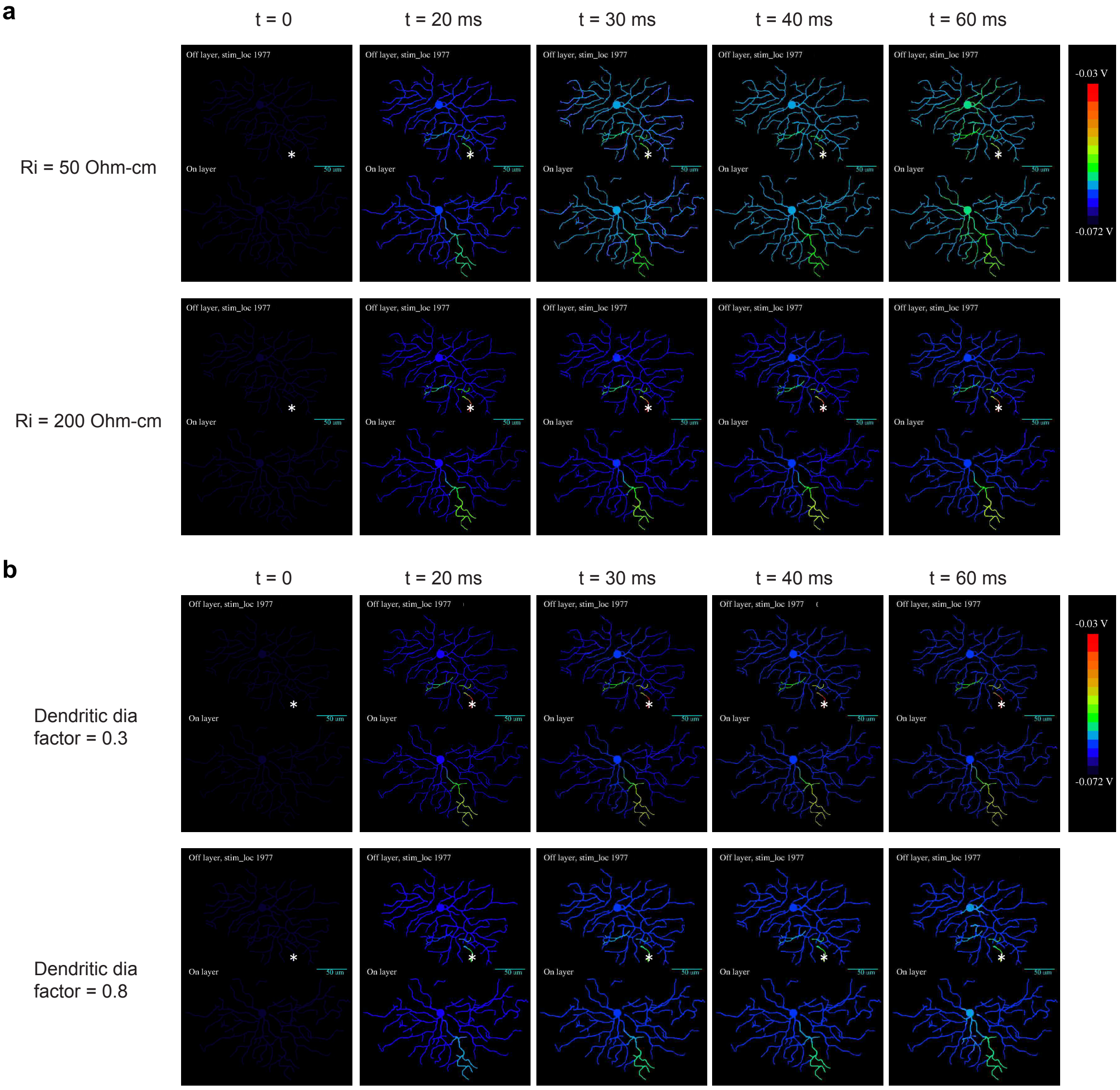


**Supplementary Figure 3. Heat maps of pDSGC membrane potentials with brackets of different parameters. Related to Fig. 6.**

**a,** Simulation model with different axial dendritic resistances. See also Fig. 6f for Ri = 100 Ohm-cm.

**b,** Simulation model with different dendritic diameter factors. See also Fig. 6f for dia factor = 0.5.

#### Supplementary Figure 4

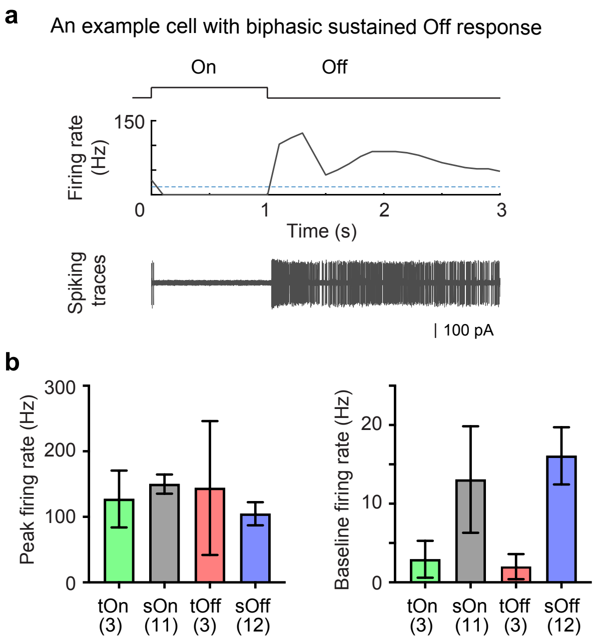


**Supplementary Figure 4. Firing patterns of alpha ganglion cells. Related to Fig. 8.**

**a,** Firing rate plot and spiking traces of an example cell with biphasic sustained Off response. The horizontal blue dash line in the firing rate plot indicates baseline firing rate when there was no visual stimulus, and the spiking traces includes 4 trials of spot responses.

**b,** Peak firing rates and baseline firing rates of four types of alpha cells. Sample sizes were represented in the plot.

#### Supplementary Figure Legends

**Supplementary Figure 1. Example spiking traces of a pDSGC represent the maintenance, extinction, and repeated induction of neural sensitization. Related to Fig. 1.**

Upper, middle and lower traces represent pDSGC spiking responses during the first, second and third trials of the induction protocol.

**Supplementary Figure 2. Heat maps of pDSGC membrane potentials after stimulation at different locations in the Off layer. Related to Fig. 6.**

The stimulated locations are indicated by * in the heat map plots. For these simulation videos, Ri = 100 Ohm-cm, dendritic dia factor = 0.5.

**Supplementary Figure 3. Heat maps of pDSGC membrane potentials with brackets of different parameters. Related to Fig. 6.**

**a,** Simulation model with different axial dendritic resistances. See also Fig. 6f for Ri = 100 Ohm-cm.

**b,** Simulation model with different dendritic diameter factors. See also Fig. 6f for dia factor = 0.5.

**Supplementary Figure 4. Firing patterns of alpha ganglion cells. Related to Fig. 8.**

**a,** Firing rate plot and spiking traces of an example cell with biphasic sustained Off response. The horizontal blue dash line in the firing rate plot indicates baseline firing rate when there was no visual stimulus, and the spiking traces includes 4 trials of spot responses.

**b,** Peak firing rates and baseline firing rates of four types of alpha cells. Sample sizes were represented in the plot.

**Supplementary Movie S1. Simulation model mimicking the electronic spread between pDSGC Off and On layers. Related to Fig. 6.**

The heat map representing the membrane potential of pDSGC after stimulation in the Off layer. The stimulated location #1977 is indicated by *. Ri = 100 Ohm-cm, dendritic dia factor = 0.5.
